# A human multilineage gut organoid model for Parkinson disease

**DOI:** 10.64898/2025.12.16.694313

**Authors:** Mahya Hosseini Bondarabadi, Minqian Xu, Jim de Leeuw, Sofie Slingerland, Sygrid van der Zee, Iris E. C. Sommer, Hermie J. M. Harmsen, Teus van Laar, Sven C. D. van IJzendoorn

**Affiliations:** Department of Biomedical Sciences, University of Groningen, University Medical Center Groningen, Groningen, the Netherlands; Department of Neurology, University of Groningen, University Medical Center Groningen, Groningen, the Netherlands; Department of Psychiatry, University of Groningen, University Medical Center Groningen, Groningen, the Netherlands; Department of Medical Microbiology and Infection Prevention, University of Groningen, University Medical Center Groningen, Groningen, the Netherlands

**Author notes:** Corresponding author S.C.D. van IJzendoorn, Department of Biomedical Sciences, University of Groningen, University Medical Center Groningen, Antonius Deusinglaan 1, 9713 AV, Groningen, the Netherlands.

## Abstract

Emerging evidence links gut dysfunction to Parkinson disease (PD) pathogenesis, yet human models to study gut-related mechanisms are lacking. We developed a human intestinal organoid model incorporating PD-relevant cell types. Using control and PD patient-derived pluripotent stem cells, we generated intestinal epithelial organoids and vagal neural crest cells, then co-cultured them into assembloids. The resultant structures featured lumen-forming polarized epithelial monolayers with enteroendocrine cells, contractile subepithelial myofibroblast-like layers, and neuroglial networks containing dopaminergic and cholinergic enteric neurons. Notably, assembloids from a PD patient carrying the GBA1-E326K variant exhibited progressive α-synuclein accumulation in non-enteroendocrine epithelial cells. This model recapitulates key gut architecture and PD-associated phenotypes, offering a physiologically relevant platform for mechanistic studies and therapeutic discovery targeting gut-brain pathways in PD.

## Introduction

Parkinson disease (PD) is the second fastest-growing neurodegenerative disorder globally and manifests through both motor deficits and a wide array of non-motor symptoms, including neuropsychiatric disturbances, autonomic dysfunction, sleep abnormalities, and gastrointestinal (GI) complications [1,2]. Although the etiology of PD remains incompletely understood, converging evidence suggests that interactions between genetic susceptibility and environmental factors promote the misfolding and aggregation of α-synuclein, a 140-amino-acid presynaptic protein central to PD and dementia with Lewy bodies pathology [3]. Under physiological conditions, alpha-synuclein modulates synaptic vesicle trafficking, but in pathological states its aggregation initiates neurotoxic cascades that contribute to progressive neuronal dysfunction and degeneration [4].

Historically viewed as a primary central nervous system disorder, PD is now conceptualized as a disease that can originate in the periphery [1]. In 2000, Braak and colleagues proposed that alpha-synuclein pathology may first arise in the enteric nervous system (ENS), following environmental triggers that breach the intestinal epithelial barrier and seed α-synuclein aggregation in enteric neurons. This pathology could then spread to the brainstem via retrograde vagal transport. In support, α-synuclein aggregates have been detected by immunohistochemistry in ENS components before motor symptom onset [5] and GI symptoms are nearly ubiquitous among PD patients, with constipation often predating motor symptom onset by many years [6].

Animal studies have shown that gut-introduced α-synuclein preformed fibrils can propagate to the brain [7–9]. These models typically rely on overexpression systems or exogenous fibril administration, limiting their physiological relevance. Demonstrations of movement of α- synuclein initiated in enteroendocrine cells to vagal neurons in mouse co-culture systems and in a transgenic SNCA^A53T^ model [10] further support a gut-originating propagation route.

This “gut-to-brain” concept has since been refined to include two proposed phenotypes, body-first and brain-first PD, that differ in their temporal sequence of pathology. Increasing evidence further indicates that specific genetic risk variants may predispose individuals to one subtype over the other [1]. Of these, risk variants in the *GBA1* gene have been correlated with gut-first PD [11].

Mounting evidence points to widespread alterations along the GI tract in PD. The role of the gut is multifactorial and likely includes a combination of genetic risk variants (most notably variants in the *GBA1* gene), environmental exposures, and multiple cell types in the gut. These include i) bacteria in the gut lumen, ii) intestinal epithelial cells (IECs) that maintain gut wall barrier and enteroendocrine cells (EECs), iii) cells in the submucosa that make up the enteric nervous system (ENS) including neurons and glial cells, and iv) other cells that regulate epithelial function such as subepithelial myofibroblasts. Abnormalities have been reported across epithelial, neuronal, glial, microbial, and immune compartments. Intestinal epithelial barrier dysfunction in PD subjects [12–14] and altered tight-junction protein profiles in patient tissue [14,15] suggest early mucosal compromise. In parallel, gut microbiome dysbiosis and heightened mucosal immune activation are consistently observed in both patients and animal models [16–18], regularly co-occurring with epithelial barrier disruption. ENS pathology has also been documented: parasympathetic denervation of the intestine [19], ganglionic degeneration in the submucosal and myenteric plexuses [20], and altered enteric glial cell signatures, including increased GFAP expression [21] and localized gliosis [22]. Despite these insights, the mechanisms that initiate α-synuclein aggregation within the gut in early stages of the disease remain unresolved.

A major drawback in studying these mechanisms is the absence of human experimental models that integrate epithelial, neuronal, glial, and genetic components of the gut while expressing endogenous α-synuclein. Such systems are required to delineate the spontaneous onset of α-synuclein aggregation, identify stimuli that trigger pathogenic processes, and determine the temporal sequence of early gut events in the pathophysiology of PD.

Pluripotent stem cell–derived organoids provide a tractable platform for *in vitro* reconstruction of human tissue architecture. Building on previous work integrating vagal neural crest cells into intestinal organoids [23], we have generated a control and PD patient derived human intestinal organoid system composed of an epithelial compartment and an embedded enteric nervous system. We conducted a structural and molecular characterization of these organoids, focusing on features directly relevant to PD, including epithelial tight-junction organization, the presence of enteric neurons and glial cells, and the expression of endogenous α-synuclein, within a single human system.

## Results

### Generation of pluripotent stem cell-derived human intestinal epithelial organoids

Human pluripotent stem cells (PSCs) were differentiated into human intestinal organoids (HIOs). We adapted a previously established stepwise differentiation protocol [24,25], incorporating minor modifications to improve robustness across PSC lines. PSC-derived embryoid bodies were differentiated over six days into mid-hind endodermal aggregates characterized by high expression of the caudal type homeobox (CDX)2 protein (Figure 1C). Following treatment with the glycogen synthase kinase 3α/β inhibitor CHIR99021 and fibroblast growth factor (FGF)4, the endodermal aggregates exhibited budding morphologies, and were subsequently embedded in Matrigel for four weeks to mature into human intestinal organoids (HIOs) with the characteristics of epithelial architecture (Figure 1D). HIOs remained stable for four to six passages without notable loss of morphology or viability. HIOs, embedded in Matrigel, consisted of a monolayer of CDX2⁺ (Figure 1E) and CDH1/cadherin⁺ intestinal epithelial cells (IECs) exhibiting apical brush border polarity, confirmed by immunolabeling for villin, a microvillus marker (Figure 1F). In addition to IECs, HIOs contained mucus-secreting cells (Figure 1G) and α-synuclein-expressing chromogranin A (CGA)-positive enteroendocrine cells (EECs) (Figure 1I).

**Fig. 1.**
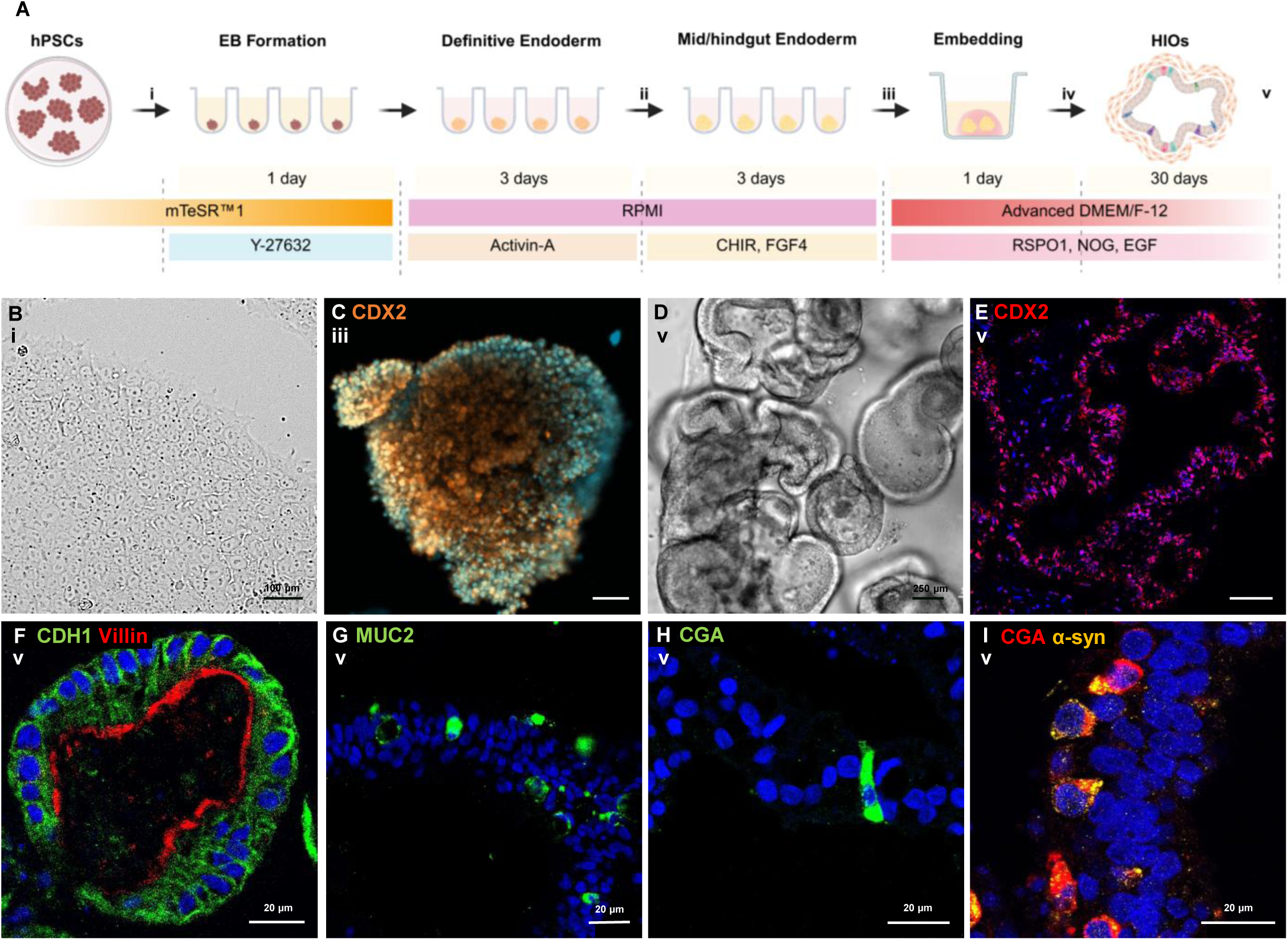
Generation and characterization of human intestinal organoids (HIOs) from pluripotent stem cells. (A) Schematic of directed differentiation of PSCs into intestinal organoids. (B–I) Images corresponding to the stages labeled i–v in the schematic. (B) Pluripotent stem cells before dissociation. (C) An embryoid body after 3 days of exposure to mid–hindgut endodermal medium acquires intestinal identity and extensively expresses the CDX2 marker. (D) Mature intestinal organoids. (E) Immunolabeled image of HIOs after 30 days in culture expressing CDX2. (F) Intestinal organoids contain enterocytes with brush borders localized on the luminal surface. (G) MUC2^+^ cells show the presence of Goblet cells within the HIOs. (H) Enteroendocrine cells within the organoids. (I) Expression of α-synuclein (α-syn) in enteroendocrine cells. Scale bars = 50 µm unless otherwise indicated. DAPI is shown in blue.

### Generation of PSC-derived vagal neural crest cells

Vagal neural crest cells (vNCC) are the primary source of precursor cells that - *in vivo* - migrate from the neural tube to the developing gut to form the enteric nervous system (ENS). Human PSC were differentiated to vNCCs following a previously established protocol [23] (Methods). The differentiation scheme is depicted in Figure 2A. The neurospheres after posteriorization were transferred to fibronectin-coated plates, where migratory vNCCs emerged within 2–3 days (Figure 2D). Immunofluorescence staining showed that the majority of delaminated cells expressed the neural crest marker SRY-Box transcription factor (SOX)10, whereas the non-migratory neurosphere core was enriched for Paired box (PAX)6, consistent with central neural fate as shown in Figure 2E. Upon passaging, vNCCs retained SOX10 expression and expressed βIII-tubulin (TUBB3) and human neuronal proteins C and D (HuC/D; also known as ELAVL3/ELAVL4), which are pan-neuronal markers (Figure 2F-H), of which HuC/D is typically expressed in more differentiated neurons.

**Fig 2.**
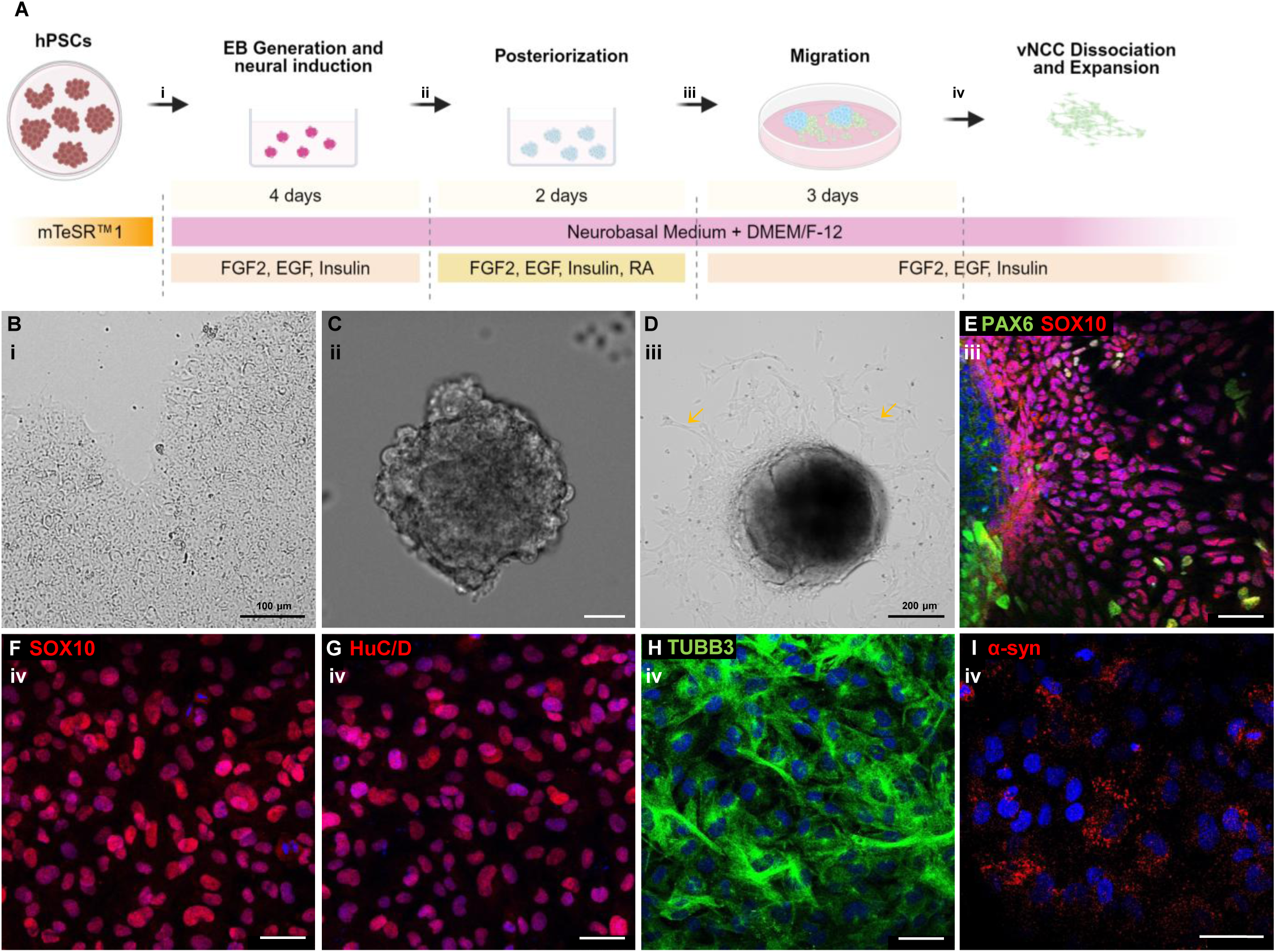
Generation and characterization of Vagal Neural Crest Cells (vNCCs) from pluripotent stem cells. (A) Schematic presentation of the directed differentiation of PSCs into vagal neural crest cells. (B–I) Distinct stages of differentiation corresponding to the schematic labels. (D) vagal NCCs delaminate on fibronectin-coated surface. (E) Migratory vNCCs derived from neurospheres abundantly express the early neural crest–specifying marker SOX10 but are largely negative for PAX6. (F) Vagal NCCs after passaging, stained for SOX10. (G, H) Vagal NCCs stained for neuronal markers. (I) Expression of α-synuclein in vagal NCCs. Scale bars = 50 µm unless otherwise indicated. Blue = DAPI.

### Assembly of HIOs and vNCCs generates a multi-lineage structured gut tissue

To establish gut organoids incorporating enteric neurons, HIO fragments were co-cultured with vNCCs as outlined in Figure 3A for 4 to 8 weeks (Supplementary Figure 1). During the first 15–30 days, cultures underwent notable structural reorganization, resulting in dense, free-floating tissues approximately 3–5 mm in diameter (Figure 3B). After 30 days in Matrigel, the co-culture produced highly organized structures in which lumen-forming polarized IECs were consistently surrounded by spatially oriented layers of smooth muscle actin (ACTA2)⁺ cells (Figure 3C). These one-to-three cell thick layers of ACTA2^+^ cells overlapped with the subepithelial deposition of extracellular matrix proteins such as collagen, as evidenced by Masson’s trichrome staining (Figure 3D). These cells therefore are highly reminiscent of intestinal subepithelial myofibroblasts.

**Fig 3.**
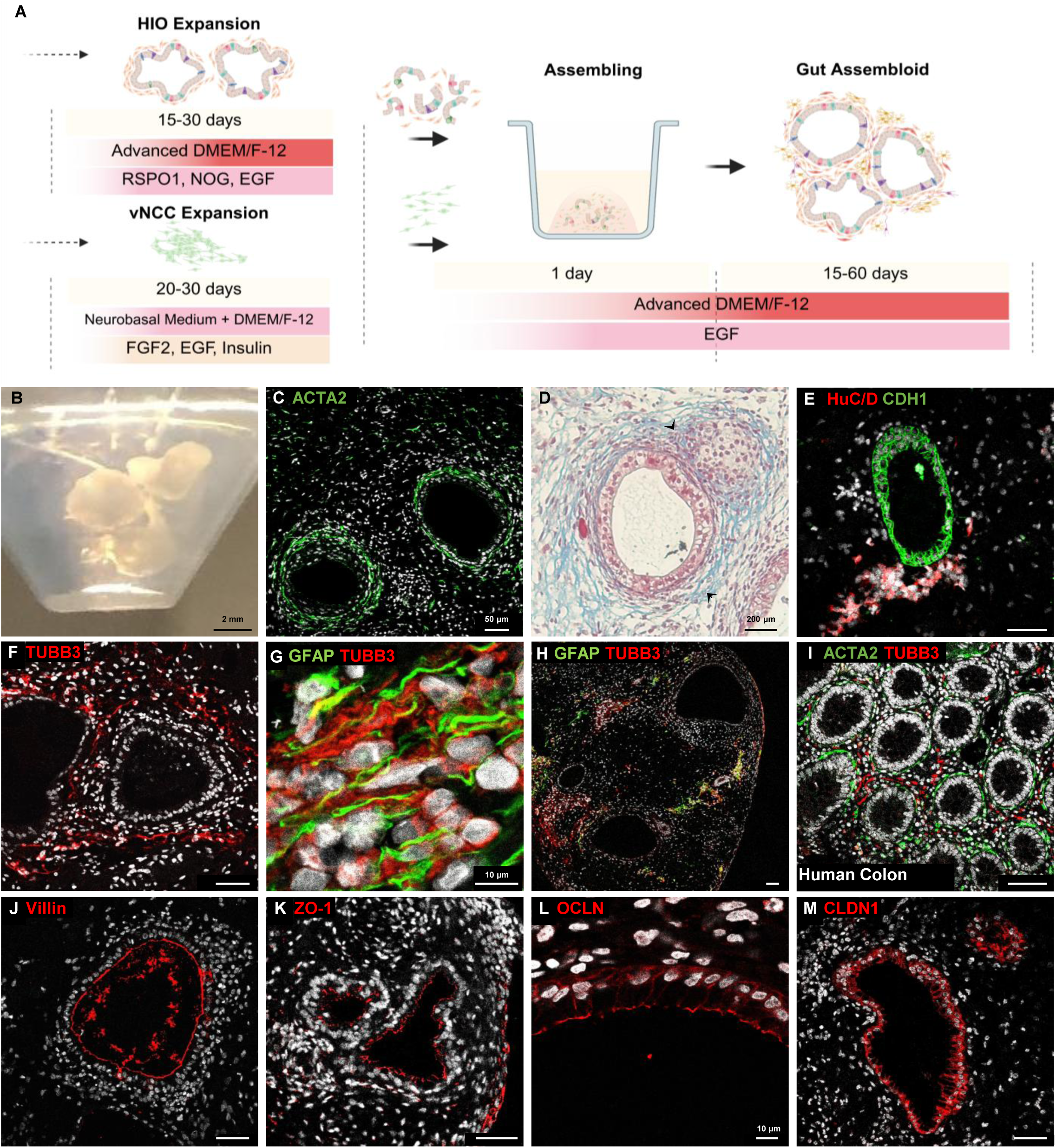
Generation and characterization of PSC-derived gut assembloids after 30 days. (A) Overview of the process for coculturing vagal neural crest cells with intestinal organoid fragments. (B) Image of three 30-day-old gut assembloids. (C) Immunostaining for smooth muscle actin (ACTA2) at the outermost layer of the intestinal epithelium. (D) Masson’s trichrome staining revealing ECM deposition beneath the epithelial layer. (E) Co-staining for HuC/D and CDH1 showing neurons located adjacent to the epithelial layer. (F) Neuronal cells distributed throughout the mesenchymal compartment of the gut assembloids. (G) High-magnification imaging showing the close spatial association between GFAP⁺ glial processes and TUBB3⁺ neuronal cells. (H) Co-staining of GFAP and TUBB3 indicating the presence of glial and neuronal populations forming plexus-like structures within the co-culture. (I) Reference human colon tissue showing the distribution of ACTA2⁺ smooth muscle cells and TUBB3⁺ neurons. (J) Villin expression at the luminal/apical surface of the intestinal epithelium. (K–M) Expression of the tight-junction proteins ZO-1 (K), Occludin (L), and Claudin-1 (M) in IECs of the gut assembloid. Nuclei are counterstained with DAPI (white). Scale bars = 50 μm unless otherwise indicated.

Beneath the ACTA2^+^ cell layer, discrete dense clusters of HuC/D⁺ and TUBB3⁺ neurons were observed (Figure 3E and F), intertwined with glial fibrillary acidic protein (GFAP)-positive glial cells (Figure 3G). Western blot analysis revealed the expression of multiple GFAP isoforms (Supplementary Figure 2). The organization of these neuro-glial networks closely resembled the submucosal plexus *in vivo* (Figure 3H). Comparative analysis with human colon tissue demonstrated striking structural similarities, particularly in the arrangement of ACTA2^+^ cells and the submucosal plexus (Figure 3I). These neuro-glial cell clusters were frequently observed in close proximity to the epithelial layer. Polarized IECs displayed tight junctions at the apical surface, as shown by immunostaining for ZO-1 (a cytoplasmic junction-associated protein) and occludin and claudin-1 (transmembrane proteins) (Figure 3 J-M). These so-called gut assembloids, following the nomenclature consensus [26], were reproducible in replicates and independent experiments.

### IECs and vNCCs are co-responsible for the differentiation of contractile ACTA2^+^ cells

The presence of ACTA2⁺ cells in the assembloids was unexpected, as these cells are of mesenchymal origin. Notably, in PSC-derived HIO cultures lacking vNCCs, only few ACTA2⁺ cells were observed with no apparent spatial organization (Supplementary Figure 3A). By contrast, in HIOs-vNCCs assembloids, many ACTA2⁺ cells were present in highly organized cell layers (Figure 3C), indicating that their differentiation and spatial patterning was stimulated by the presence of vNCCs.

Increasing the number of vNCCs within Matrigel-embedded HIOs-vNCCs assembloids resulted in a marked increase - up to 70% - in tissue contraction and suspension of the tissue (Figure 4A and B). Such contraction was also observed - albeit to a lesser extent - in vNCCs monocultures (Figure 4B and Supplementary Figure 4A) but not observed in HIO monocultures (Figure 4A and B). In the compacted assembloids, occasional colocalization of ACTA2⁺ cells with TUBB3⁺ neurons was observed (Figure 4C and D), suggesting neuro-mesenchymal interactions. The observed compaction aligns with the established role of active subepithelial myofibroblasts in extracellular matrix remodeling and gel contraction.

**Fig 4.**
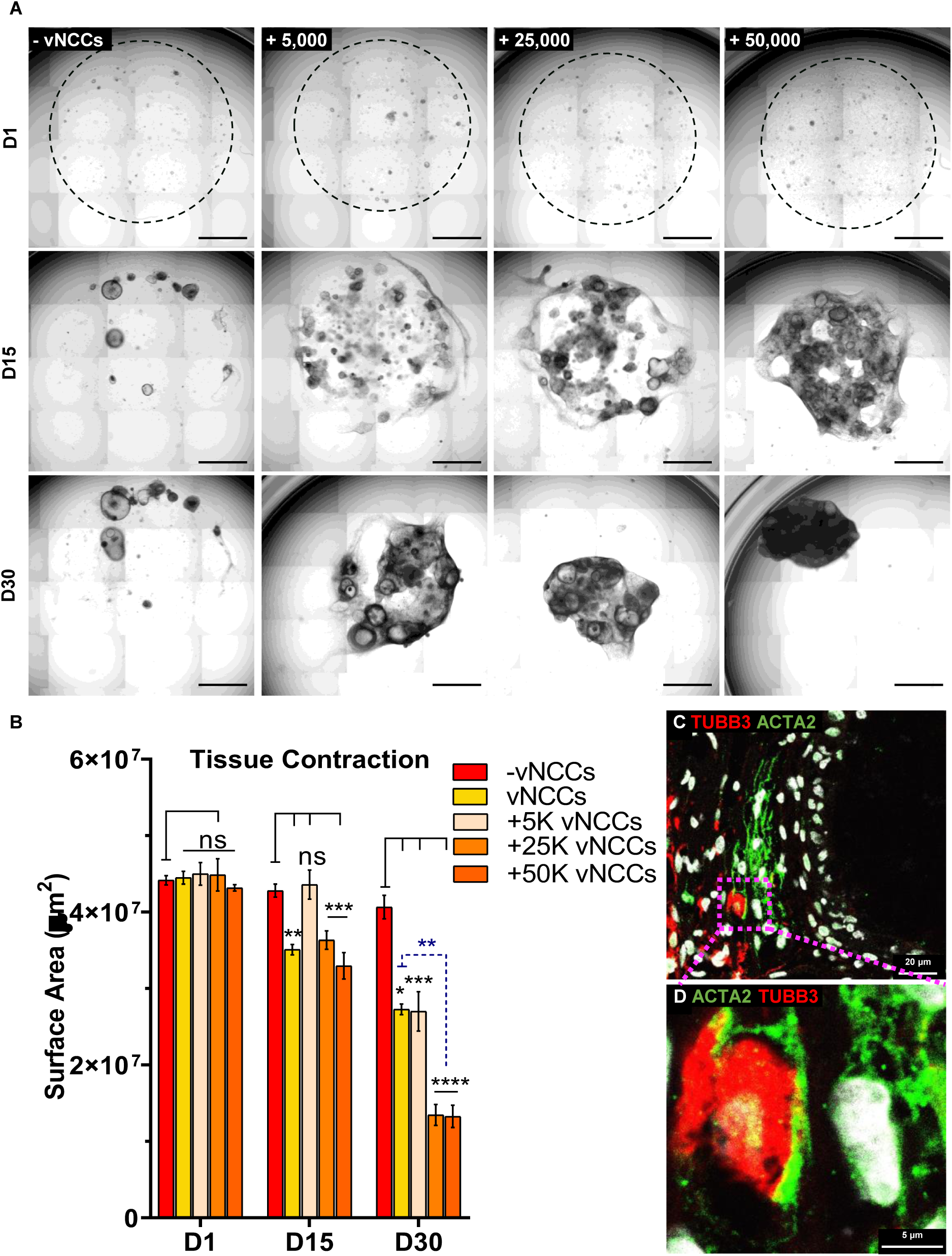
Restructuring of the co-culture of HIOs and vNCCs during time. (A) Images show HIOs cultured alone or in the presence of 5 × 10³, 25 × 10³, and 50 × 10³ vagal neural crest cells. Each row illustrates the time-course reorganization of the entire structure over days 1, 15, and 30, driven by the cross-talk between HIOs and vNCCs. Scale bars = 1000 µm. (B) Quantification of the surface area of Matrigel beads containing HIOs and vNCCs at different time points demonstrates gradual contraction of the structure in co-cultures. (C) TUBB3⁺ neuronal cells are colocalized with ACTA2⁺ cells in the underlying subepithelial tissue. (D) Magnified image of panel C. Nuclei are counterstained with DAPI (white).

Importantly, no collagen deposition and no tissue contraction were observed when vNCCs were co-cultured with liver cell spheroids (Supplementary Figure 4A-C), indicating that the differentiation and spatial organization of contractile ACTA2⁺ cells required both vNCCs and HIOs. Together with the profound spatial orientation of ACTA2^+^ cells around HIOs, our findings suggest functional cross-talk between HIOs and vNCC-derived cells to achieve mesenchymal differentiation and patterned organization.

### PSC-derived gut assembloids include PD-relevant dopaminergic and cholinergic neuronal subtypes

TUBB3+ and HuC/D+ neuronal cells in the HIOs-vNCCs assembloids were further characterized by immunolabeling for various neuronal subtypes. The assembloids contained tyrosine hydroxylase (TH)-positive neurons (Figure 5A), which are present in the *in vivo* gut ENS as a dopaminergic subset and the loss of which has been associated with altered colonic motility, a well-known non-motor PD symptom [27]. The assembloids also contained choline acetyltransferase (ChAT)-positive neurons (Figure 5B), which in the *in vivo* gut ENS are the predominant cholinergic neurons driving excitatory neurotransmission via acetylcholine [28]. Figure 5C illustrates how these neurons project neurite-like outgrowths to the intestinal epithelial layer. As shown in Figure 5D, neurons in the assembloids also expressed ubiquitin carboxyl-terminal hydrolase L1 (UCHL1), which is important for maintaining protein homeostasis and neuron health, and the loss of which caused age-related neurodegenerative changes in the ENS [29]. Taken together, these findings demonstrate that the HIOs-vNCCs assembloids supported robust neurogenesis from vNCCs and that PD-relevant neuronal subtypes were present.

**Fig. 5.**
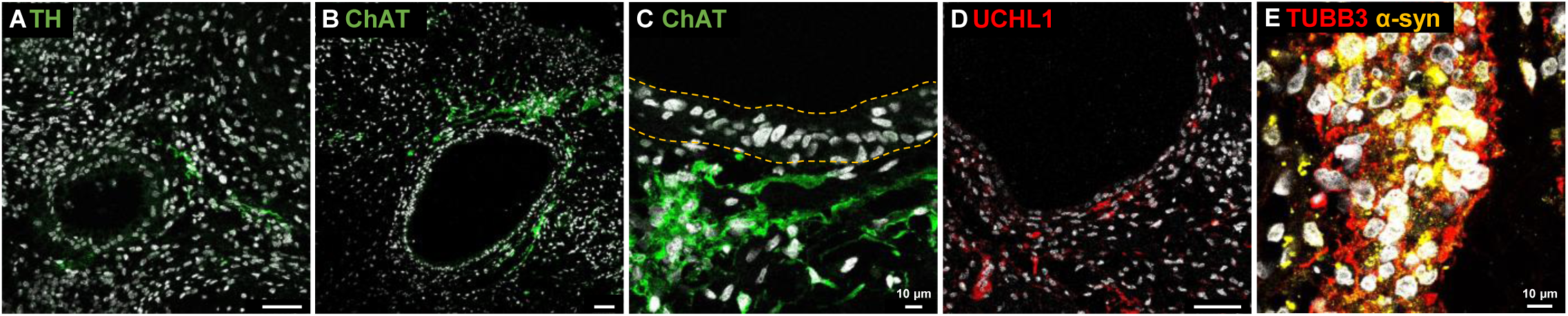
Gut assembloids exhibit abundant enteric neuronal populations. (A) TH⁺ dopaminergic neurons are consistently detected within gut assembloids. (B) Cholinergic neurons are distributed around the epithelial structures. (C) ChAT^+^ neurons extended projections toward the epithelial layer (dashed line). (D) UCHL1⁺ neurons are detected in gut assembloids. (E) TUBB3⁺ neuronal cells highly express α-synuclein. Scale bars = 50 µm unless otherwise indicated. Nuclei are counterstained with DAPI (white).

### Endogenous expression of α-synuclein in the gut assembloid model

To assess the relevance of the gut assembloid model for PD, we examined endogenous α-synuclein expression in human intestinal organoids (HIOs), vagal neural crest cells (vNCCs), and the thereof generated assembloids. Detection was performed using the monoclonal 15G7 antibody that targets amino acids 116–131 in the C-terminus of human alpha-synuclein (Supplementary Table 1).

The specificity of this anti-α-synuclein antibody was validated in a genetically engineered human Flp-In™ T-REx™ 293 cell line stably expressing wild-type α-synuclein fused to DsRed2 under a tetracycline-inducible promoter (pcDNA5/FRT/TO-aSYN(WT)-DsRed2; gift from dr. Eduardo Mattos and dr. Kampinga (University Medical Center Groningen, the Netherlands). Expression of α-synuclein was observed by immunofluorescence microscopy and western blot only in tetracycline-induced cells (Supplementary Figure 5A and B).

In HIOs, α-synuclein was primarily detected in CGA^+^ enteroendocrine cells (Figure 1I), consistent with *in vivo* findings [10], and only occasional presence of α-synuclein was observed in non-EEC intestinal epithelial cells. Expression was also observed in vNCC-derived cells (Figure 2I) and in neuro-glial cells within the assembloid (Figure 5E). Across all cell types, α-synuclein appeared as punctate cytoplasmic structures, matching its reported *in vivo* distribution.

These results confirm that α-synuclein is expressed in the appropriate cell types within the gut organoid model.

### PD patient-derived gut assembloids show progressive accumulation of α-synuclein in non-enteroendocrine epithelial cells

Having established the gut assembloid model, we next generated assembloids from induced pluripotent stem cells (iPSCs) derived from a Dutch PD patient. This patient was heterozygous for the E326K variant in the *GBA1* gene. *GBA1* variants have been associated with body/gut-first PD [11]. The pluripotency of the iPSC line was confirmed by the expression of pluripotency markers and the ability of the iPSC to differentiate toward all embryonic germ layers (Supplementary Figure 6A-C and E). The presence of the *GBA1*-E326K variant was confirmed by Sanger sequencing and a normal karyotype was confirmed (Supplementary Figure 6D). PD patient-derived iPSCs were differentiated to HIOs and vNCCs, which were morphologically indistinguishable from their respective controls (Supplementary Figure 6B).

Co-culture of patient-derived HIOs and vNCCs produced PD^GBA1-E326K^ assembloids with polarized CDH1⁺ epithelial monolayers with apical tight junctions, surrounded by ACTA2⁺ myofibroblast-like cells and spatially organized neuro-glial clusters (Supplementary Figure 6H-K), similar to control assembloids (*cf.*, Figure 3).

To further investigate potential differences in α-synuclein expression patterns between control and patient-derived gut assembloids, multiple sections from both cultures were examined by immunostaining. Neuro-glial cell populations did not exhibit visibly altered α-synuclein expression in both control and patient-derived gut assembloids in two different time points (Figure 6B, G, D, and I). However, in contrast to controls (Figure 6A and F), IECs in PD^GBA1-E326K^ assembloids showed a significant accumulation of α-synuclein (Figure 6C and H). Upon prolonged culture for up to two months, IECs in PD^GBA1-E326K^ assembloids exhibited larger α-synuclein aggregate-like structures which appeared most prominent at the basal side of the cells as illustrated in Figure 6J. These results were further validated by using a different antibody (the 211 clone) directed against amino acids 121-125 of human α-synuclein (supplementary Figure 7). Occasionally we observed α-synuclein-expressing TUBB3^+^ neuronal cells in very close proximity to the α-synuclein-expressing IECs (Supplementary Figure 8A). Importantly, these α-synuclein-expressing IECs were negative for CGA (Supplementary Figure 8B), a marker for EECs which are known to express α-synuclein (*cf*., Figure 2I; [10]).

**Fig. 6.**
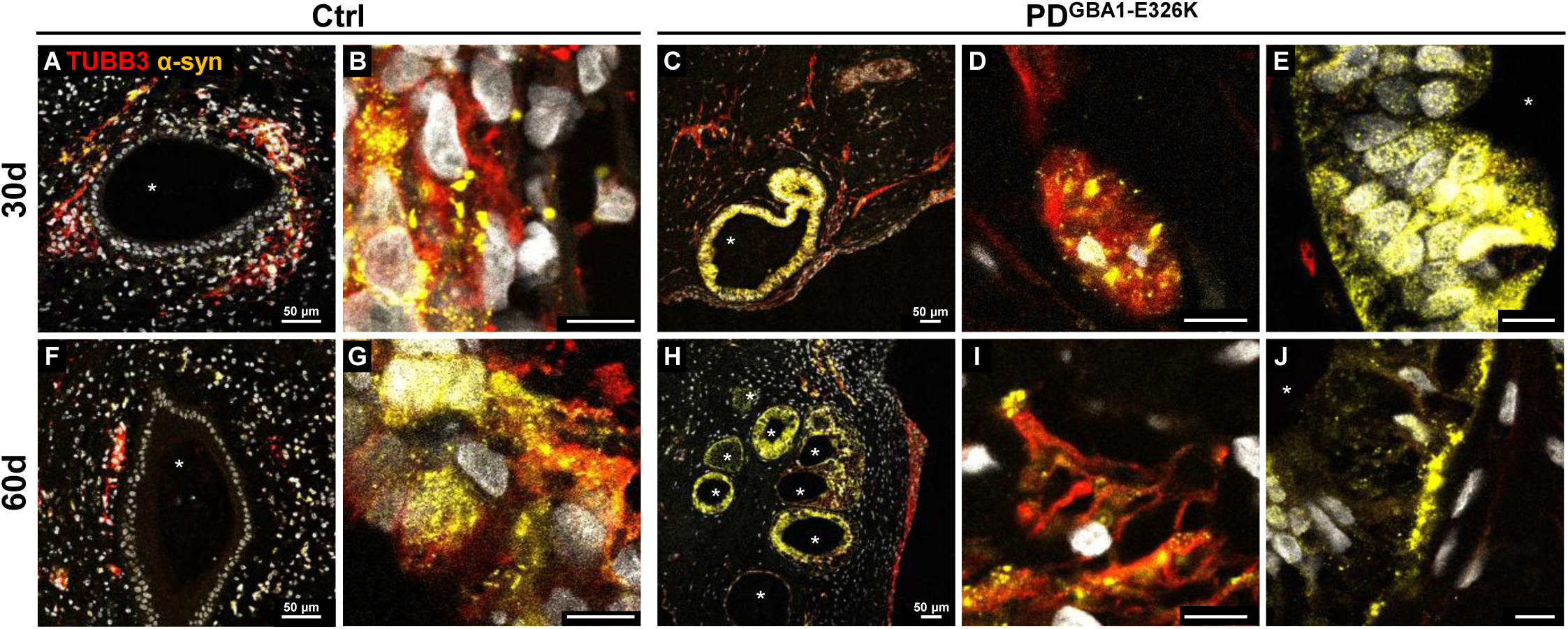
Detection of different patterns of α-synuclein within the IEC layer of gut assembloids derived from control and PD-iPSCs. (A–B and F–G) Co-staining for TUBB3 and α-synuclein in control gut assembloids; (C–E and H–J) staining in PD GBA1-E326K gut assembloids. (A, F) IEC layers of control assembloids show no detectable α-synuclein. (B, G) α-synuclein is strongly localized within TUBB3⁺ neuronal processes in control gut assembloids. (C, H) In PD^GBA1-E326K^ assembloids, IECs are positive for α-synuclein in both 30-day and 60-day cultures. (D, I) The pattern of α-synuclein expression in TUBB3⁺ neurons remains similar between 30-day and 60-day cultures. (E, J) In PD^GBA1-E326K^ assembloids, the diffuse α-synuclein signal observed in IECs at day 30 becomes more punctate after 60 days of culture. DAPI is shown in white. The luminal region of the HIO indicated by *. Scale bars = 10 µm unless otherwise indicated.

## Discussion

So far, most research on the pathophysiology of Parkinson disease (PD) has been focused on the brain. Studies with patients and experimental mouse models have convincingly demonstrated that the gut plays a role in the etiology and/or progression of PD. However, thus far, no experimental human model existed that combine these proposed essential players in the PD gut.

### Development of a gut organoid model for Parkinson disease

To enable gut-focused investigations of PD pathophysiology, we generated millimeter-sized human pluripotent stem cell (PSC)-derived gut assembloids visible to the naked eye. These gut assembloids integrate lumen-forming, polarized intestinal epithelial cells (IECs) monolayers that also contain mucin-producing cells and enteroendocrine cells (EECs). These are surrounded by perpendicularly orientated 1-3 cell thick layers of contractile subepithelial myofibroblasts and populations of dopaminergic and cholinergic neuronal subtypes entangled with glial cells in a submucosal plexus-like organization. Endogenous expression of the α-synuclein protein, a key protein in the pathogenesis of PD, was readily detected in the neuro-glial cells as well in EECs located in the epithelial layer. In these cells, α-synuclein appeared as discrete punctate structures in the cytoplasm. The α-synuclein protein was not typically observed in non-EECs IECs, in agreement with the notion that IECs do not transcribe the α-synuclein-encoding *SNCA* gene. We have thus generated a human gut assembloid that contains the most Parkinson disease-relevant cell types in an *in vivo*-like configuration.

The assembloids unexpectedly showed highly organized and orientated layers of ACTA2^+^ cells beneath the epithelial layer, overlapping with similarly orientated layers of collagen and therefore highly suggestive for intestinal subepithelial myofibroblasts. Such extensive organization of HIOs and NCC-derived cells was thus far only observed following their co-transplantation into recipient mice [30]. The origin of these ACTA2^+^ cells in the *in vitro* assembloids is not clear. Subepithelial myofibroblasts have been suggested to originate from neural crest cells and/or from the serosal mesothelium [31,32]. The emergence of smooth muscle fibroblasts and neuronal cells has also been observed in iPSC-derived HIOs *in vitro* when cultured in a microfluidic chip [33] and after treatment with retinoic acid [34] or epiregulin [35] (*i.e.*, all in the absence of vNCCs). It has been proposed that these culture conditions may have stimulated differentiation of residual neuronal/mesenchymal progenitors following endoderm/hindgut differentiation [34]. ACTA2^+^ cells were not reported in the pioneering study by the Wells lab, where also HIOs and vNCCs were co-cultured to generate assembloids [23], which may be due to protocol differences. Our results indicate that in our assembloids vNCCs and HIOs cooperatively promoted the differentiation and spatial organization of these cells. Significant shrinkage and compaction of assembloids correlated with the number of vNCCs used to generate the assembloids. Because active myofibroblast generate traction forces that can shrink collagen gels [36], we therefore consider ACTA2^+^ myofibroblast-like cells in the assembloid to display contractile activity.

### Parkinson disease patient-derived gut assembloids show aberrant presence and progressive accumulation of α-synuclein in intestinal epithelial cells compared to assembloids from controls

We generated induced pluripotent stem cell (iPSC) lines from a Dutch PD patient carrying a heterozygous GBA1 c.1093G>A (p.E326K) variant. GBA1 variants, including E326K, are associated with approximately a two-fold increased risk of PD compared to non-carriers, often linked to gut-first PD, earlier onset, and faster progression of motor and non-motor symptoms [11]. Experimental models have implicated GBA1-E326K in α-synuclein pathology: ectopic expression of GBA1-E326K in human cell lines promotes α-synuclein aggregation [37], and injection of preformed α-synuclein fibrils into the gut of Gba1-E326K knock-in mice accelerated Lewy body accumulation in the hippocampal dentate gyrus and worsened non-motor symptoms [38]. Gba1-E326 expression thus promoted pathogenic α-synuclein transmission and intensified disease pathology in a mouse model. However, the effects of the endogenously expressed heterozygous GBA1-E326K variant on α-synuclein has not been studied before.

Presentation of α-synuclein staining did not show differences between neuro-glial cells in control and PD^GBA1-E326K^ assembloids. We did observe a significant increase in the presence of α-synuclein in IECs in PD^GBA1-E326K^ assembloids compared to controls. In these cells, α-synuclein appeared as small punctate structures spread throughout the cytoplasm in 30 days-old assembloids, but upon prolonged culture for up to two months exhibited a larger, aggregate-like appearance at the basal side of the IECs. These α-synuclein-positive IECs did not express CGA, which is a marker for EEC located within the IEC layer and known to express α-synuclein [10]. Inspection of publicly available single-cell RNA-seq datasets of mouse and human intestinal epithelium indicates that SNCA transcripts are largely restricted to enteroendocrine cell clusters, with negligible or no detectable expression in other epithelial lineages [10]. Nonetheless, multiple studies have reported reactivity to α-synuclein antibodies in non-EEC IECs in gut biopsies of PD patients, inflammatory bowel disease patients and controls [12,39–43], with one study reporting positive staining of α-synuclein in 42% of PD gut biopsies analyzed [43]. This phenomenon, however, has received little attention and often has been dismissed as being non-specific staining because IECs do not transcribe the *SNCA* gene.

We can hypothesize that α-synuclein detected in IECs of PD^GBA1-E326K^ assembloids may have originated from subjacent enteric neurons. Enteric neurons can transfer free or exosome-encapsulated α-synuclein between cells [44,45]. The α-synuclein protein in IECs was most prominent at the basal side facing submucosal cell types, and TUBB3+ neurons expressing the α-synuclein protein could be observed adjacent to α-synuclein-positive IECs. The increase in α-synuclein protein in IECs carrying the GBA1-E326K variant when compared to controls may reflect impaired clearance of transmitted α-synuclein or stimulated aggregation of α-synuclein, consistent with previous findings of α-synuclein aggregation observed in human cell lines overexpressing GBA1-E326K [37]. Future studies are needed to test this hypothesis, whether this observation is specific for the GBA1-E326K variant, and whether α-synuclein accumulation in IECs affects IECs function.

Taken together, we generated human intestinal assembloids from control and PD patient-derived pluripotent stem cells, integrating epithelial, mesenchymal and enteric neural components. These structures mimic gut architecture, express endogenous α-synuclein and show PD phenotypes, including progressive α-synuclein accumulation. This gut assembloid model provides a unique opportunity to investigate early PD gut pathology, explore patient-specific susceptibility, and evaluate therapeutic interventions in a preclinical setting.

## Methods

### Human pluripotent cell culture

Induced pluripotent stem cells were generated from peripheral blood mononuclear cells (PBMCs) from a patient diagnosed with Parkinson disease (PD) and carrying a heterozygous GBA1-E326K variant. PBMCs were cultured for 10–12 days with medium change every other day by medium 1 containing StemPro-34 SFM (Thermo Fisher, 10639011) supplemented with GlutaMAX, penicillin/streptomycin, SCF, FLT3L, IL-3, IL-6, and erythropoietin (PeproTech). Reprogramming was performed at the iPSC-CRISPR Facility of the University Medical Center Groningen (Groningen, The Netherlands) using the Sendai virus-based CytoTune-iPS 2.0 kit (Thermo Fisher, A16517) per the manufacturer’s protocol. On transduction day, cells were maintained in medium 1, then cultured for 3 days in medium 1 enriched with sodium butyrate (Sigma, B5887) and ascorbic acid [Sigma, A4544] under hypoxia. On day 3, cells were transferred to Geltrex-coated (Thermo Fisher, A1413202) plates with inactivated MEFs. From days 4–6, cytokines were removed. From day 7, cultures were gradually transitioned to E6 medium (Thermo Fisher, A1516401) Containing FGF2. Then, the cells were cultured in E8 medium (Thermo Fisher, A1517001). iPSC colonies with typical morphology were manually picked between days 18–25 for expansion and characterization.

Pluripotent stem cells were maintained under feeder-free conditions on hESC-qualified Matrigel (Corning, 354277) in mTeSR1 medium (StemCell Technologies, 85850) at 37 °C with 5% CO₂. Cultures were passaged weekly using ReLeSR (StemCell Technologies, 100-0484) following the manufacturer’s instructions at a 1:10 ratio for ESCs and 1:6 for iPSCs.

### Differentiation of pluripotent stem cells to human intestinal organoids

PSCs were dissociated using Accutase (StemCell Technologies, 07922), and 2,500–3,000 cells were seeded into low-attachment 96-well round-bottom plates (ThermoFisher, 174925) in mTeSR1 (StemCell Technologies, 85850) supplemented with 10 µM Y-27632 (Tocris, 1254) overnight. To prevent the spheroid loss, a white background was used to improve the contrast.

Spheroids were transferred to RPMI 1640 containing 1× non-essential amino acids (Gibco, 11140050), 100 U/mL penicillin/streptomycin (Gibco, 15140122), and 100 ng/mL Activin A (Peprotech, 120-14P-500UG). Fetal bovine serum (Gibco, 16141002) was added at 0%, 0.2%, and 2% on days 1–3, respectively. From days 4 to 6, the cultures were maintained in the same basal medium supplemented with 2% FBS, 500 ng/mL FGF4 (Peprotech, 100-31), and 3 µM CHIR99021 (Tocris, 4423).

Ten to twelve mid/hindgut endoderm spheroids were embedded in Matrigel (Corning, 354230) and after polymerization of Matrigel, the culture was maintained in intestinal growth medium including Advanced DMEM/F12 (Gibco, 12634010), supplemented with 1X B27 (Gibco, 17504001), 2 mM GlutaMAX (Gibco, 35050038), 15 mM HEPES (Gibco, 15630080), 100 U/mL penicillin/streptomycin, 1 µM CHIR99021, 100 ng/mL NOGGIN (Peprotech, 120-10C-01M), 100 ng/mL EGF (Peprotech, AF-100-15-500UG), and 500 ng/mL R-SPONDIN (Peprotech, 120-38-01M). Medium was refreshed every 4–7 days. Organoids were passaged between days 14–20 after embedding, and harvested for co-culture after 3–4 passages.

### Differentiation of pluripotent stem cells to vagal neural crest cells

Human PSC colonies were mechanically disrupted by introducing random scratches with a P10 pipette tip, followed by enzymatic detachment using 500 U/mL Collagenase I (Gibco, 17018029) in mTeSR1 for 20–30 minutes under visual monitoring. Loosened fragments were collected by gentle pipetting, centrifuged at 450 × g for 5 minutes, transferred to low-attachment culture dishes (CytoOne, CC7672-3359), in neural induction medium composed of a 1:1 mixture of DMEM/F12 (Gibco, 21331020) and Neurobasal medium (Gibco, 21103049), supplemented with 0.5× B27, 0.5× N2 (Gibco, 17502001), 5 µg/mL insulin (Invitrogen, RP-10908), 20 ng/mL FGF2 (Peprotech, 120-14P-50UG), and 20 ng/mL EGF. The medium was changed daily, and 2 µM retinoic acid (Sigma, R2625) was added on days 4 and 5 for posteriorization.

On day 6, free-floating neurospheres were plated onto fibronectin-coated surfaces (3 µg/cm²; R&D Systems, 1918-FN) and cultured in neural induction medium without retinoic acid for an additional 3–4 days. Migratory cells emerging from neurospheres were detached by brief Accutase treatment (90 seconds), passed through a cell strainer (StemCell Technologies, 27215) to exclude larger clumps, and seeded onto fresh fibronectin-coated plates. Cells were used for downstream applications following a few passages.

### Assembling the HIOs–vNCCs to generate assembloids

HIOs at passages 3–4 and vNCCs at passages 2–3 were used. HIOs were mechanically dissociated into tissue fragments following retrieval from the matrix using cold DMEM/F12. For standard innervated HIO generation, 5 × 10⁴ vNCCs were combined with HIO fragments in proportions adjusted according to total tissue yield. The cell–tissue mixture was centrifuged at 450 × g for 5 minutes at 4 °C, and the resulting pellet was resuspended in 50 µL of chilled, fresh Matrigel containing 10% intestinal growth medium. After gentle pipetting to ensure even distribution without introducing bubbles, the mixture was plated into pre-warmed 24-well plates and incubated at 37 °C for 30 minutes to allow Matrigel polymerization. The co-culture was maintained in growth medium including Advanced DMEM/F12, supplemented with 1X B27, 2mM GlutaMAX, 10 mM HEPES, 100 U/mL penicillin/streptomycin, 100 ng/mL EGF for 30 to 60 days with refreshing every 4–5 days or any time considering the phenol red indicator.

For comparative experiments, HIOs were first pooled and dissociated into fragments in a single tube to ensure consistent input. Equal volumes of the resulting HIO suspension were distributed into separate tubes, and distinct numbers of vNCCs (5 × 10³, 25 × 10³, and 50 × 10³) were added to each tube prior to embedding and culture.

### Imaging for contraction

Images were acquired on day 1, 7, 15, and 30 post Matrigel embedding using the ZEISS Celldiscoverer 7 microscope. The image analysis was done by Zen3.6 (blue edition) software with the circle and spline tools to measure the surface area of the tissue.

### Immunofluorescence staining

Cells and spheroids were fixed with 4% paraformaldehyde at RT 15-30 min on a shaking platform. After permeabilization with 0.2% Triton X-100 for 15 min, samples were rinsed with PBS and blocked with 1% BSA for an hour at RT. Then the samples were exposed to the primary antibodies as listed in supplementary table 1 at 4 °C overnight. After rinsing the samples three times in PBS, they were treated with the secondary antibodies in 1% BSA and the nuclei were counterstained with DAPI (Invitrogen, D3571). Mounting Medium (Dako, S3025) was used for mounting the coverslips. The images were taken by Leica TCS SP8.

### Organoid processing and staining

Samples were fixed in 4% paraformaldehyde at 4 °C overnight. Tissues were processed in EtOH series and Xylene substitute (Merck, 1098435000) and embedded in paraffin, sectioned at 4.5 μm, and affixed to Superfrost Plus Adhesion slides (Epredia, J7800AMNZ). Paraffin sections were hydrated and underwent heat-induced epitope retrieval with Tris EDTA (pH 9.0) in microwave. Sections were rinsed with MiliQ water then exposed to the blocking buffer including 1X PBS with 1% BSA and 1% normal donkey serum (Jackson Immuno Research, 017-000-001) in PBS for 90 min at 23 °C. Primary antibodies (Supplementary Table 1) diluted in blocking buffer were applied to slides overnight at 4 °C in a humidified chamber, followed by washes and incubation with secondary antibodies and DAPI in blocking buffer at 23 °C for 60 min. Slides were mounted and images were obtained using a Leica TCS SP8. Images were processed using LAS X software.

Masson’s Trichrome staining was performed on FFPE sections following rehydration. Slides were mordanted for 15 min in preheated Bouin’s solution at 58 °C, stained with Weigert’s iron hematoxylin for 5 min, then briefly differentiated in acid–EtOH, and rinsed. Sections were then incubated in Biebrich Scarlet–Acid Fuchsin (5 min), followed by phosphomolybdic acid (2 min) and Aniline Blue (5 min). After a brief acetic acid rinse, slides were dehydrated, cleared in xylene, and mounted. Images were acquired using an Olympus BX5 microscope.

### qRT-PCR analysis

Quantitative PCR for pluripotency markers was performed on a LightCycler® 480 using the SsoAdvanced™ Universal Probes Supermix and PrimePCR™ Expression Probe Assays (Bio-Rad). For the assessment of mesodermal marker expression and confirmation of Sendai virus transgene clearance, total RNA was isolated using TRIzol reagent (Invitrogen). cDNA was synthesized using a RevertAid RT Reverse Transcription Kit. Quantitative RT-PCR was then performed using iTaq Universal SYBR Green Supermix (Bio-Rad). Reactions were run on a CFX384 Real-Time System (Bio-Rad). Gene expression levels were calculated using the ΔΔCt method, normalized to GAPDH or β-actin. Samples with undetermined amplification (Cq = N/A) were assigned a Cq value of 45 (limit of detection) for visualization and considered Sendai virus-negative.

### Protein extraction and western blotting

Proteins were extracted using RIPA protein extraction buffer (10 mM Tris pH 7.5, 150 mM NaCl, EDTA 0.5 mM, Triton X-100 1%, DOC 1%, 0.1% SDS) containing protease inhibitors (Roche, 11836153001) on ice. In total, 30-50 μg of the protein lysate measured by BCA assay was loaded on a 10-15% SDS–PAGE gel and transferred to a PVDF membrane (Millipore, IPFL85R). Blots were blocked with 5% BSA, incubated with primary antibodies (1:1000) in blocking buffer overnight at 4 °C and were then probed with secondary antibodies (Supplementary Table 1) 1:5000 in 5% BSA for 1 h at room temperature on a shaking platform. Visualization was done by Odyssey® CLx Imaging System and using the Image Studio Lite Version X.

## Statistical analysis

All statistical analyses were performed using GraphPad Prism version 8.0.2 (GraphPad Software). Comparisons between two independent groups were conducted using an unpaired two-tailed Student’s t-test. Data are presented as mean ± SEM unless otherwise specified. A *P*-value < 0.05 was considered statistically significant.

## Resource availability

## Materials availability

Materials are available upon request at the corresponding author.

## Data and code availability

Data are available upon request at the corresponding author. No code was generated in this study.

## Acknowledgements

We acknowledge the support from the Brain Foundation Netherlands (grant number JP-06) and from the Hadders-deCock Foundation (grant number 2024-30). We are grateful for dr. Bart Eggen and dr. Wia Baron (University Medical Center Groningen) for sharing reagents. We thank dr. Eduardo Mattos from the lab of dr. Harrie Kampinga (University Medical Center Groningen) for sharing the Flp-In™ T-REx™ 293 cell line stably expressing wild-type α-synuclein fused to DsRed2 under a tetracycline-inducible promoter (pcDNA5/FRT/TO-aSYN(WT)-DsRed2). We thank Nieske Brouwer, Ricky Thijssen, and Klaas Sjollema (University of Groningen Imaging Center) and Mathilde Broekhuis (University of Groningen iPSC and CRISPR facility) for technical assistance. Part of the work has been performed at the UMCG Imaging and Microscopy Center (UMIC).

## Author contributions

Overall conceptualization, MH, TL, SI and SIJ; methodology, MH, JL, MX, SS, and SZ; investigation, MH, JL, MX, and SZ; resources, SS, SZ, TL; writing – original draft, MH and SIJ; writing – review and editing, all authors; supervision, SIJ, TL, HH and IS; funding acquisition, SIJ and IS.

## AI Statement

During the preparation of this work the author(s) used Copilot in order to check for text clarity. After using this tool/service, the author(s) reviewed and edited the content as needed and take(s) full responsibility for the content of the publication.

## Supplementary figure legends

**Supplementary Fig 1.**
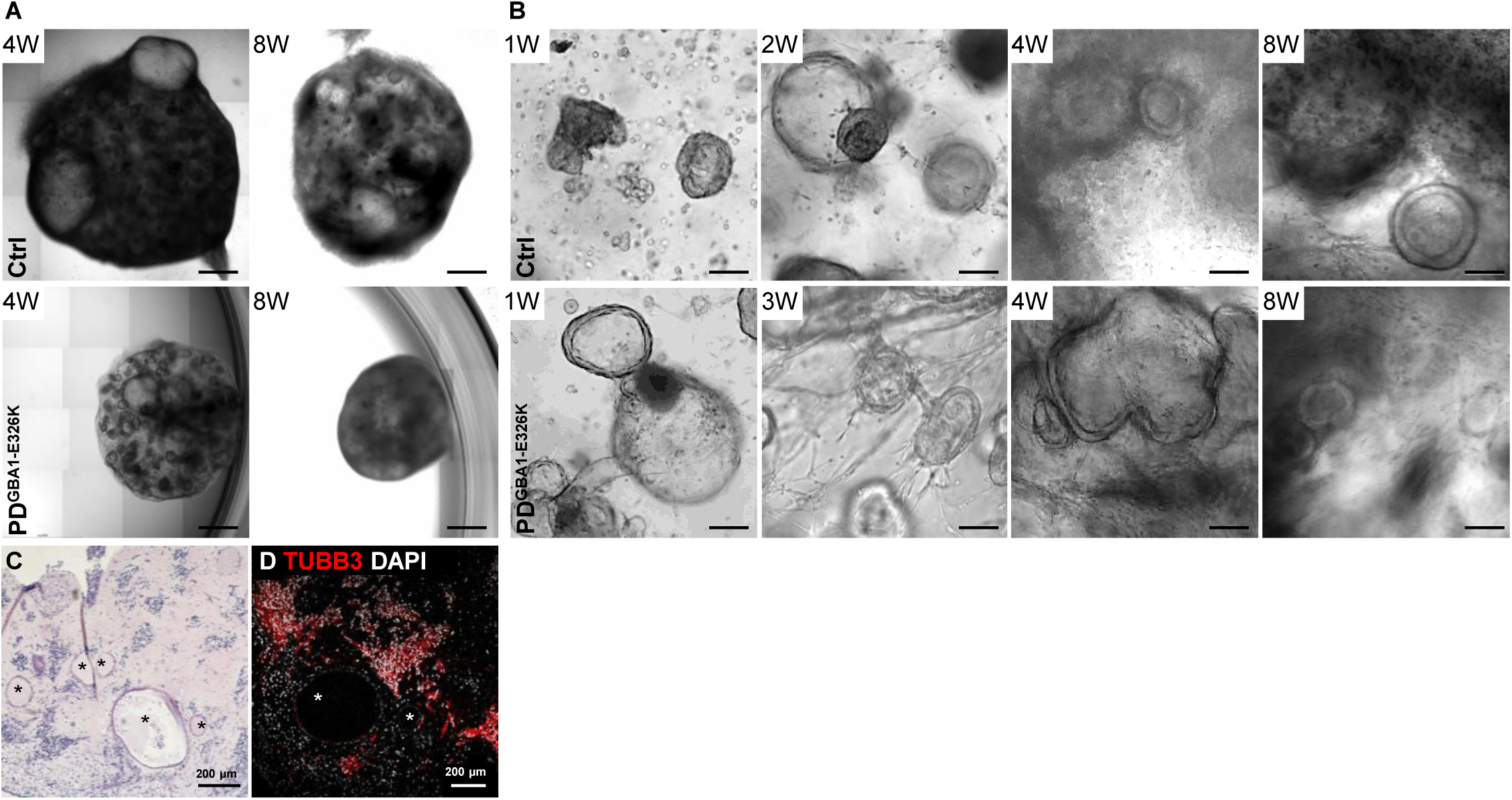
Long-term maintenance of gut assembloids in culture. (A) Morphology of ESC-derived (top row) and PD^GBA1-E326K^-derived (bottom row) HIOs-vNCCs co-cultures after 4 weeks (4W) and 8 weeks (8W). Scale bars = 1000 µm. (B) Chronological bright-field images showing morphological changes from week 1 to 8 in culture. Scale bars = 100 µm. (C) H&E-stained section of an 8-week-old gut assembloid showing preserved tissue architecture. (D) TUBB3 staining marking neuronal structures, with the luminal region of the HIOs indicated by *.

**Supplementary Fig. 2.**
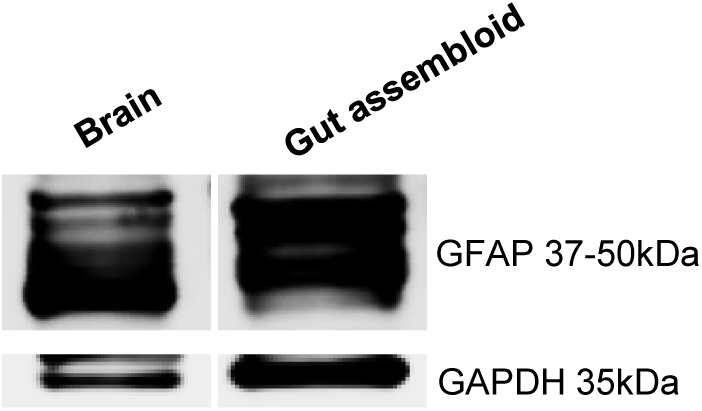
Western blot validation of GFAP isoforms in gut assembloids. Western blot analysis confirms expression of different GFAP isoforms (37–50 kDa) in gut assembloids derived from ESCs compared with brain tissue homogenate. GAPDH (35 kDa) served as a loading control.

**Supplementary Fig. 3.**
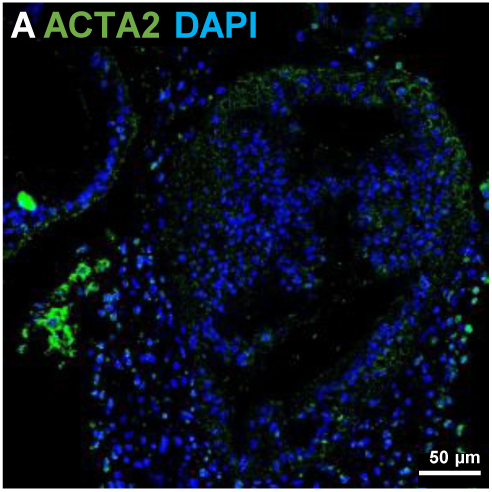
ACTA2⁺ cells are sparsely distributed within HIOs monocultures. (A) Immunostaining shows scattered ACTA2 expression in the mesenchymal compartment surrounding the intestinal epithelium.

**Supplementary Fig 4.**
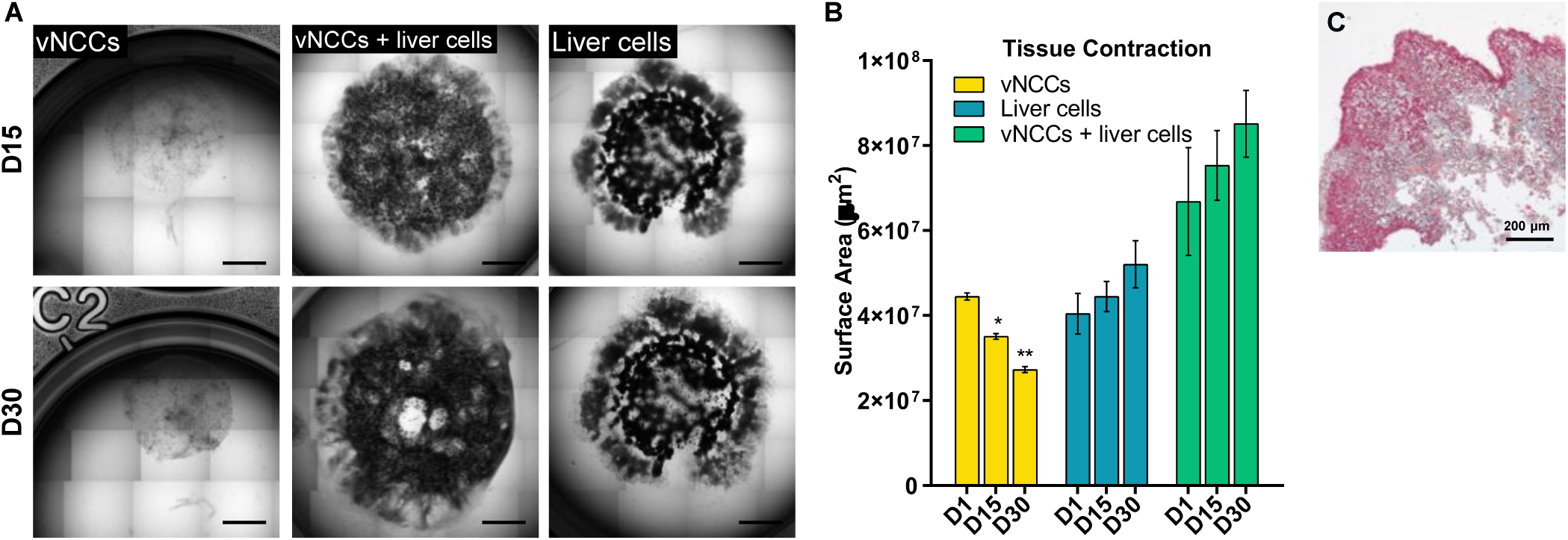
The reorganization is specific to HIOs and vNCCs cross-talk. (A) Images show the single-culture of 50 × 10³ vNCCs embedded in Matrigel beads or combined with 100 × 10³ liver cells. Each row illustrates the reorganization of the entire structure after 15 and 30 days in culture. Scale bars = 1000 µm. (B) Quantification of the surface area of Matrigel beads containing vNCCs with or without liver cells at different time points. (C) Scattered deposition of ECM in liver cells-vNCCs co-culture.

**Supplementary Fig. 5.**
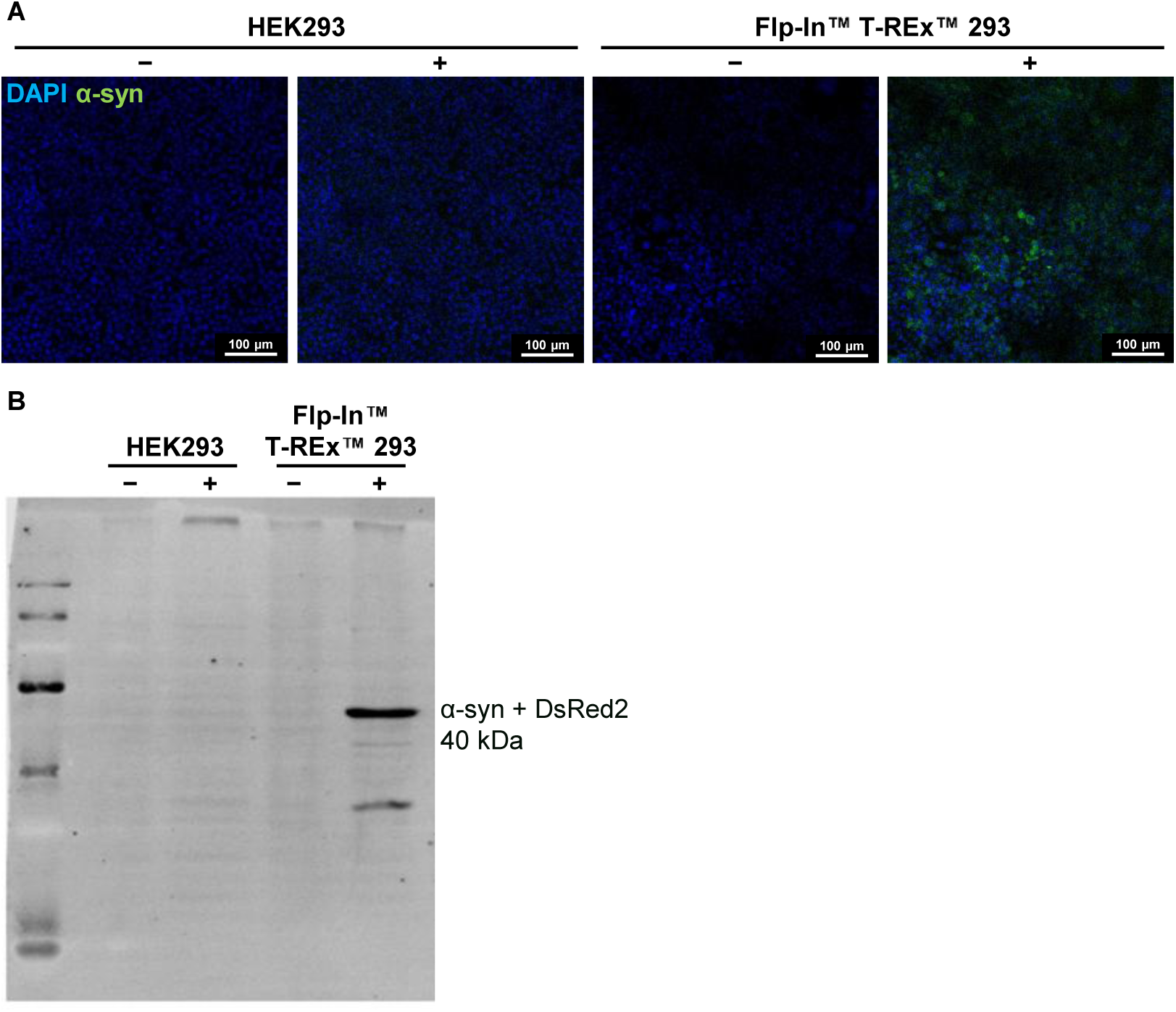
Validation of anti-α-synuclein antibody specificity. (A) Immunofluorescence analysis of HEK293 cells and genetically engineered Flp-In™ T-REx™ 293 cells stably expressing wild-type α-synuclein in the presence (+) or absence (−) of tetracycline. Nuclei are counterstained with DAPI (blue), and α-synuclein fused to DsRed2 is detected by immunolabeling using the 15G7 antibody and an anti-rat Alexa Fluor 647 secondary antibody (shown in green), independent of the DsRed2 fluorescent tag. (B) Western blot analysis confirming α-synuclein expression exclusively in tetracycline-induced Flp-In™ T-REx™ 293 cells, with no detectable signal in uninduced cells or parental HEK293 controls.

**Supplementary Fig. 6.**
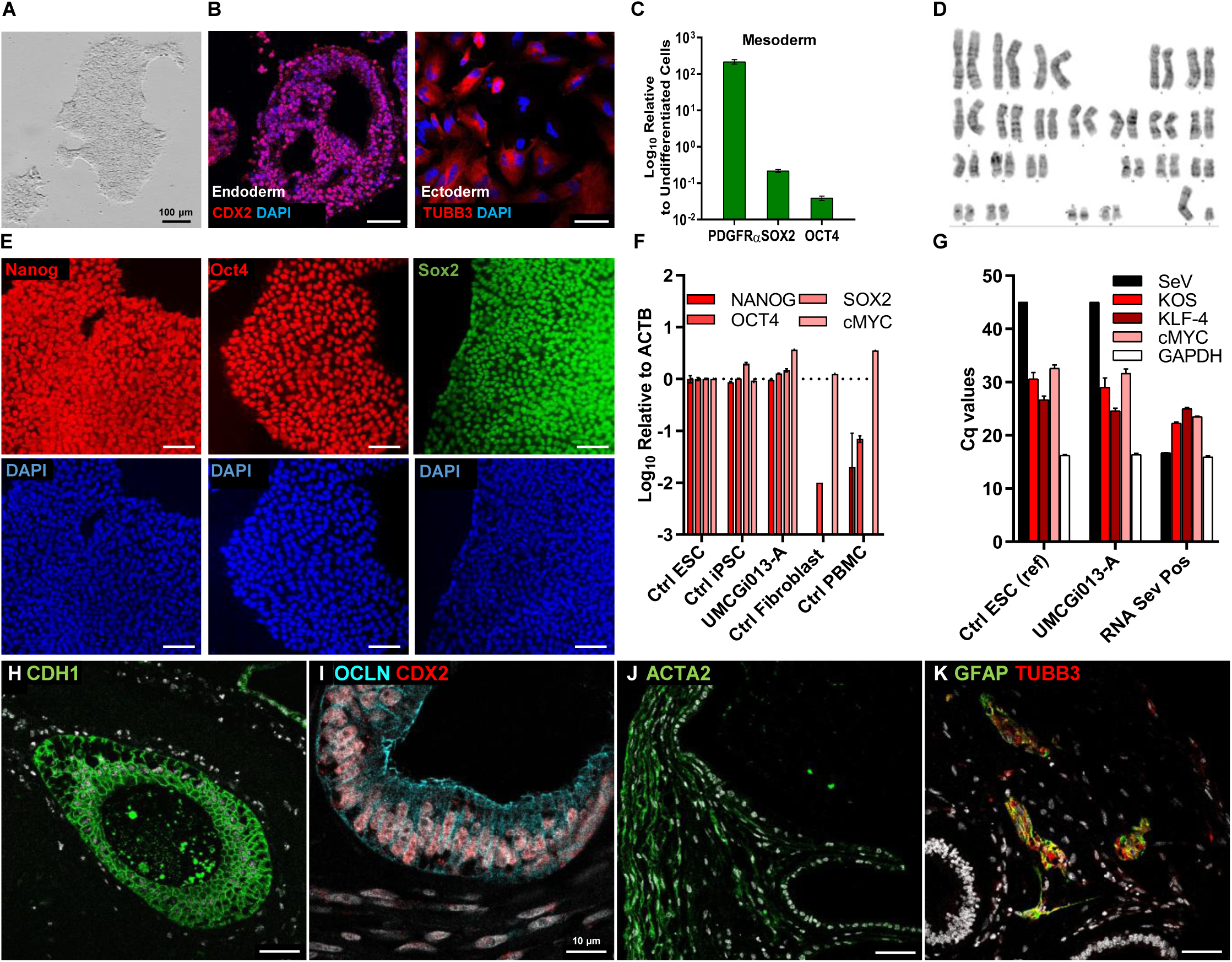
Characterization of the UMCGi013-A (GBA1-E326K) iPSC line and the gut assembloids derived from it. (A) UMCGi013-A iPSCs carrying GBA1-E326K variant exhibit normal ESC-like morphology. (B, C) The trilineage differentiation capacity of the line is confirmed by immunostaining and qRT-PCR. (D) Karyotyping shows a normal XY karyotype. (E, F) UMCGi013-A iPSCs express pluripotency markers. (G) The line is free of residual Sendai virus. (H) Gut assembloids derived from this line express the enterocyte marker CDH1. (I) Assembloids also express the intestinal marker CDX2 and the tight-junction protein Occludin, and (J) contain ACTA2⁺ cells at the subepithelial domain. (K) Immunostaining reveals submucosal-like plexus structures in gut assembloids generated from the GBA1-E326K patient line. Scale bars = 50 μm unless otherwise indicated. DAPI is shown in white in panels H–K.

**Supplementary Fig. 7.**
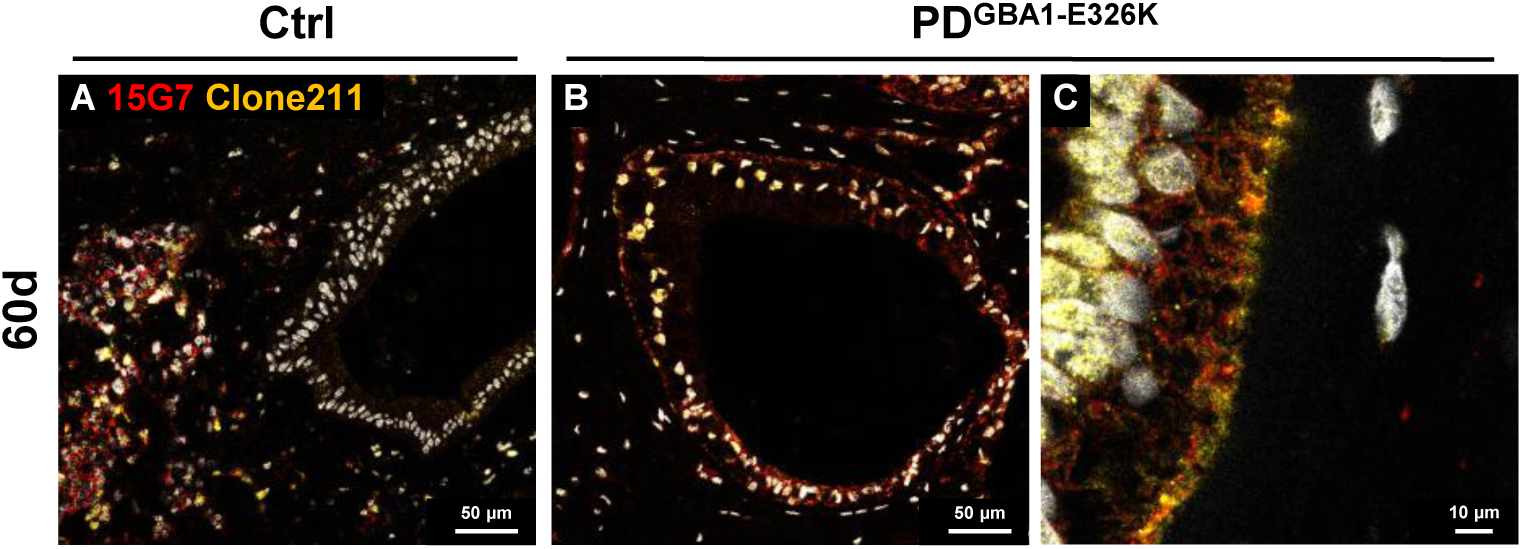
α-synuclein is detected in control and PD GBA1-E326K assembloids using epitope-distinct antibodies. Representative immunolabeled images of control (A) and PD GBA1-E326K (B, C) gut assembloids stained with an independent anti-α-synuclein antibody recognizing a distinct epitope (clone 211), confirming the presence of α-synuclein in IEC layer of the PD gut assembloid. Nuclei are counterstained with DAPI (white).

**Supplementary Fig. 8.**
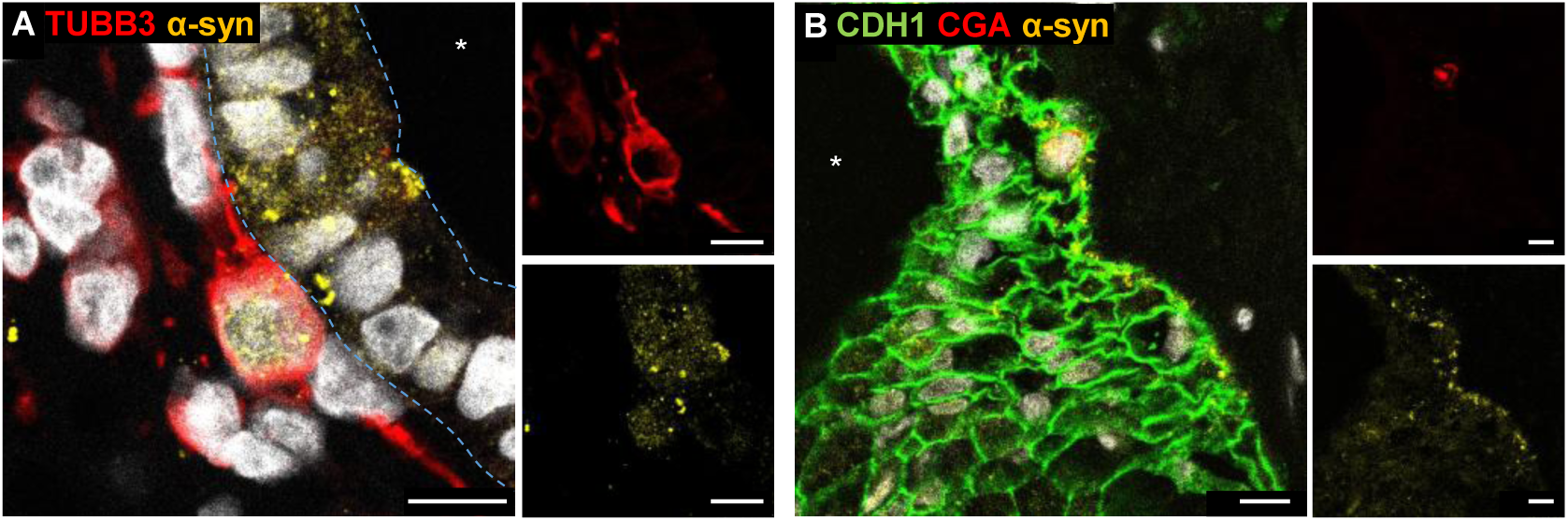
α-synuclein localization in distinct cells within gut assembloids. (A) Co-staining for TUBB3 and α-synuclein (α-syn) showing α-synuclein puncta in neuronal cells and adjacent intestinal epithelial cells. (B) Co-staining for CDH1, CGA, and α-synuclein demonstrating α-synuclein expression in non-CGA^+^ IECs. Nuclei are counterstained with DAPI (white). The luminal region of the HIO indicated by *. Scale bars = 10 µm.

## Supplementary Tables

**Supplementary Table 1.**
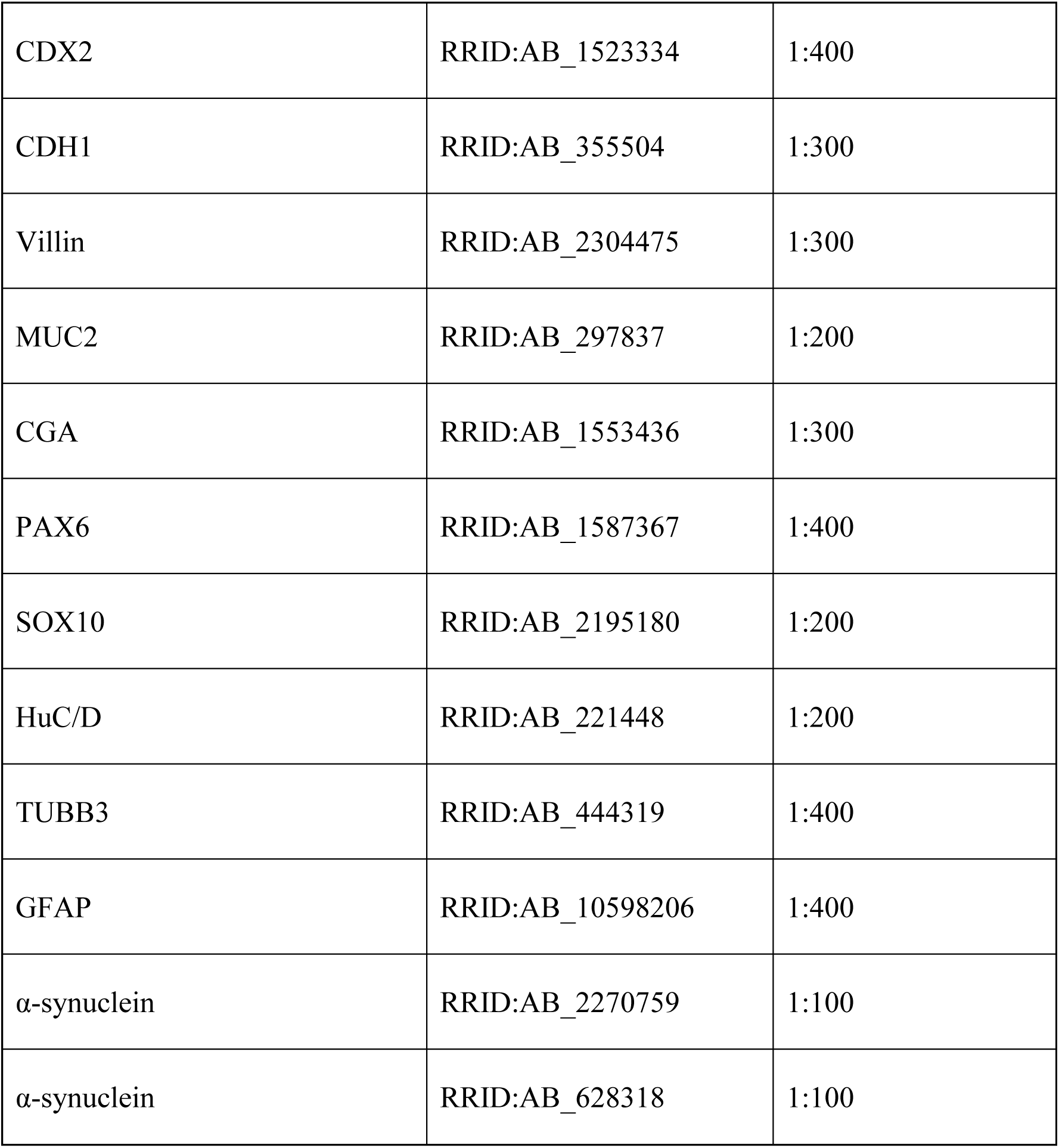
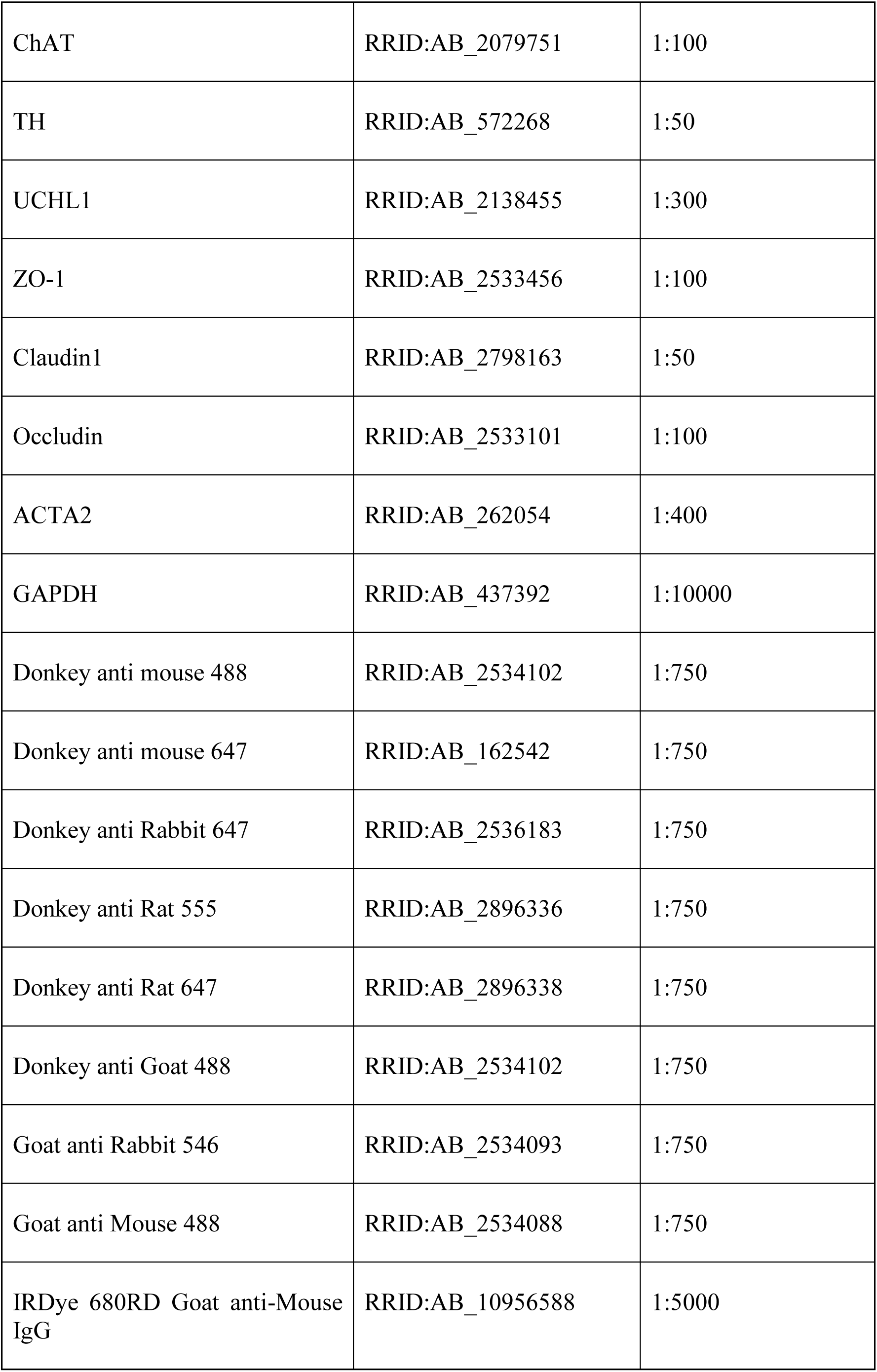
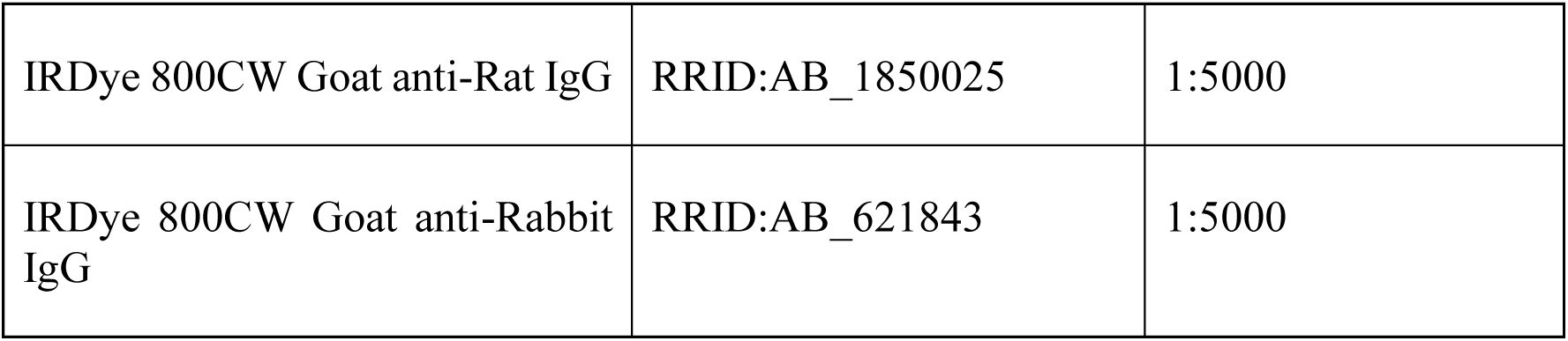
List of antibodies used.

**Supplementary Table 2.**
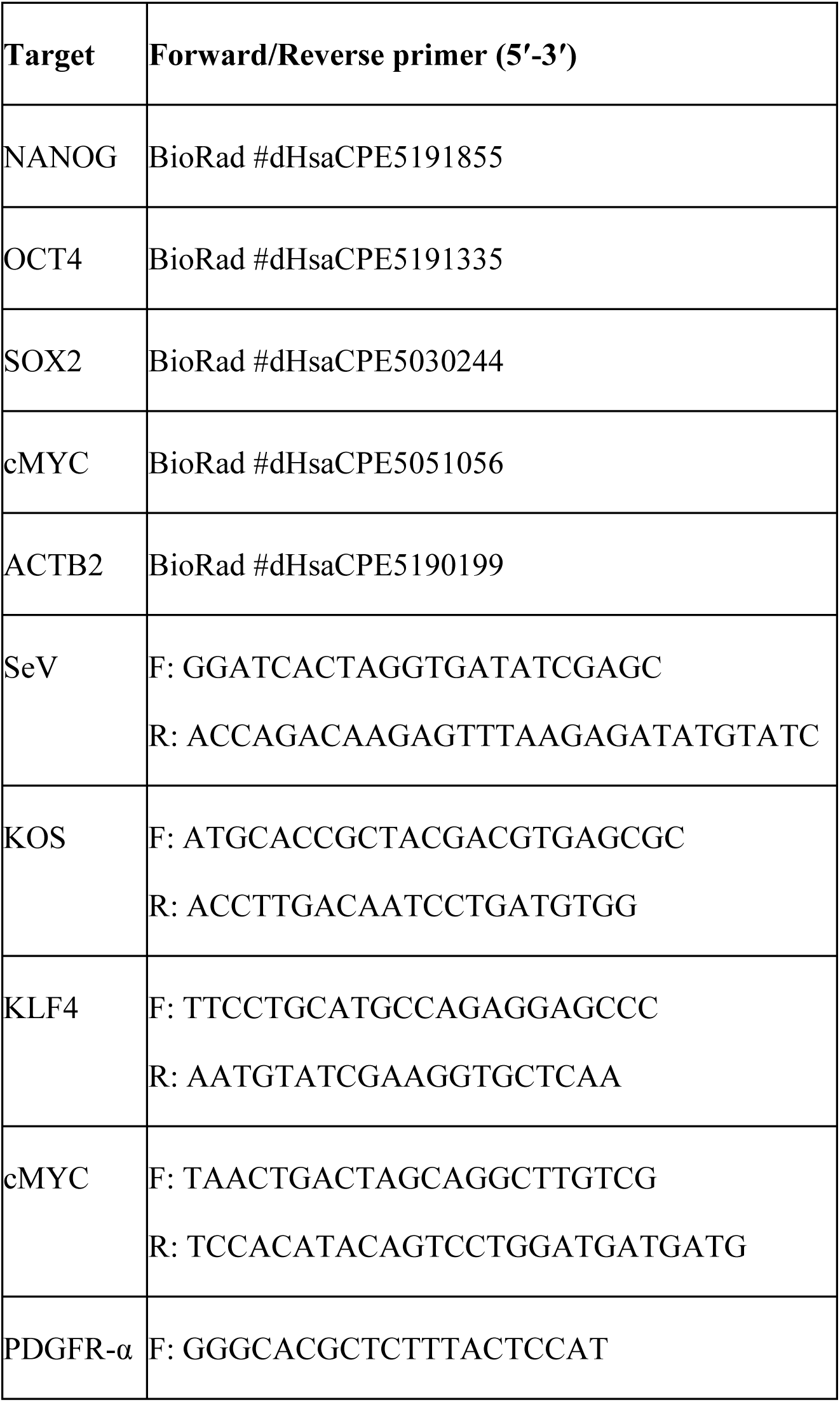
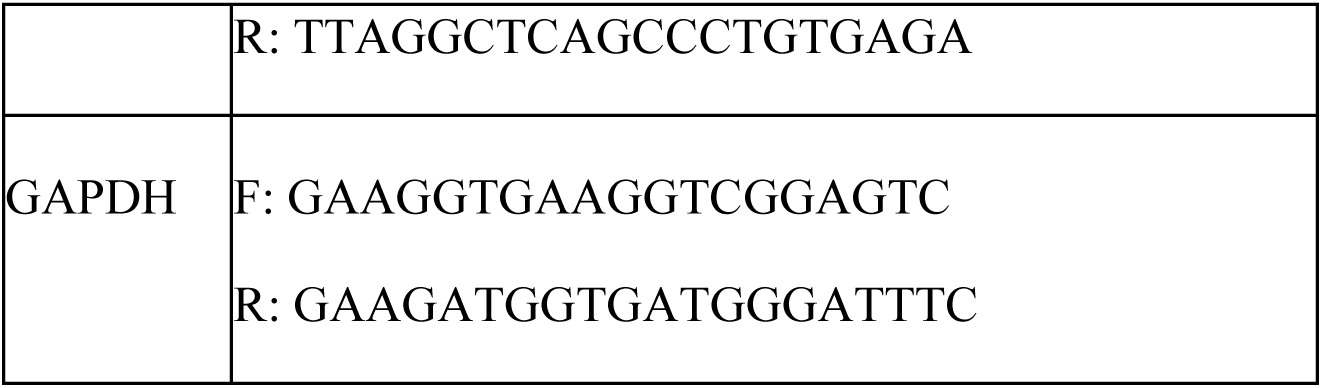
List of primers used.

